# Barcoding biological reactions with DNA-functionalized vesicles

**DOI:** 10.1101/672287

**Authors:** Justin A. Peruzzi, Miranda L. Jacobs, Timothy Q. Vu, Neha P. Kamat

## Abstract

Targeted vesicle fusion is a promising approach to selectively control interactions between vesicle compartments and would enable the initiation of biological reactions in complex aqueous environments. Here, we explore how two features of vesicle membranes, DNA tethers and phase-segregated membranes, promote fusion between specific vesicle populations. We show that membrane phase-segregation provides an energetic driver for membrane fusion that increases the efficiency of DNA-mediated fusion events. Using this system, we show that orthogonality provided by DNA tethers allows us to direct fusion and delivery of DNA cargo to specific vesicle populations. We then demonstrate that vesicle fusion between DNA-tethered vesicles can be used to initiate *in vitro* protein expression that leads to the synthesis of model soluble and membrane proteins. The ability to engineer orthogonal fusion events between DNA-tethered vesicles will provide a new strategy to control the spatio-temporal dynamics of cell-free reactions, expanding opportunities to engineer artificial cellular systems.

## Introduction

Controlling biological reactions spatially and temporally remains a significant challenge in the design of cell-mimetic systems. Cellular systems utilize extracellular vesicles, like exosomes and ectosomes, to transport biological molecules to target cells and initiate biochemical reactions.^[1,2]^ This type of chemical communication often requires that vesicles fuse with their target membrane in order to discharge their cargo into the cellular cytoplasm and initiate biological processes such as gene expression.^[3,4]^ Mimicking this process of targeted vesicle fusion with synthetic vesicles has the potential to transform a variety of technologies, ranging from drug delivery to the design of artificial cellular systems.

Specifically, the ability to engineer fusion events between distinct populations of vesicles would allow for new and complex biochemical behaviors. DNA-mediated fusion could allow for the targeted delivery of cargo to vesicle-based bioreactors, especially when vesicles reside in hard to reach environments, like the blood stream. Targeted fusion could allow for such bioreactors to be reloaded with new reagents, promoting longer durations of product synthesis. Multiple reactions could also be simultaneously executed in the same environment with limited opportunities for reaction cross-talk. Finally, complex multi-step reactions that require reagents or products to be mixed in a specific sequence in order to limit off-target product synthesis could be better engineered. While previous work has shown that lipid vesicles can be engineered to fuse in order to initiate cell-free reactions,^[5–7]^ the ability to rapidly and noninvasively control fusion between specific populations of vesicles while limiting off target fusion to others has not yet been achieved.

DNA-programmed vesicle fusion provides a promising approach to direct the targeted fusion of distinct populations of vesicles. Introduced and initially developed by the H⍰⍰k and Boxer groups, these systems use lipid-anchored DNA tethers that hybridize complementary DNA strands on different vesicle membranes.^[8,9]^ Hybridization can subsequently drive fusion between vesicles, leading to content mixing of luminal cargoes. Since this initial study, several groups have investigated methods to enhance fusion efficiency, generally by altering oligonucleotide architecture and chemistry,^[10–16]^ or have introduced methods to control the sequence of fusion events by controlling the identity of DNA tethers on different populations of vesicles.^[17]^ However, the impact of membrane biophysical features on DNA-mediated vesicle fusion dynamics and the application of DNA-mediated vesicle fusion to drive biochemical reactions have yet to be explored. An important feature of vesicle membranes that should significantly affect fusion is membrane organization. In cellular systems, localized structures known as lipid rafts are expected to play an important role in a variety of cellular processes related to membrane deformation, including exocytosis, endocytosis, signaling, and fusion.^[18–20]^ These structures are expected to be important for membrane fusion events as they create an energetic landscape which makes membrane fusion and reformation more favorable.^[21]^ Liquid-ordered domains are generally thicker than liquid-disordered domains.^[22]^ This discrepancy in membrane thickness generates both an energetic cost due to interfacial energy between phases, known as line tension, and induces local membrane curvature changes to accommodate mismatches in membrane thickness.^[23,24]^ Upon membrane fusion, the energetic cost due to the presence of lipid domains may be relieved to some degree through fusion of domains on opposing membranes.^[25]^ In synthetic membranes, domain formation can be readily engineered by choosing specific compositions of membrane components^[26]^ and has been shown to enhance the efficiency of peptide-mediated fusion.^[25,27]^ Yet, the potential of membrane domains to impact fusion events for the purpose of technological or therapeutic applications has to date, received limited investigation^[28]^ and merits further exploration.

Here, we demonstrate how features of vesicle membranes, including membrane domains and membrane hydrophobic mismatch at domain boundaries, promote efficient vesicle fusion (Figure 1 A) in DNA-tethered systems. We first investigate how biophysical features of vesicle membranes can be used to control the extent of membrane fusion. We then show that the molecular specificity of DNA-oligonucleotides functionalized to vesicle membranes enables DNA cargo to be delivered to specific vesicle populations (Figure 1 B). Lastly, we demonstrate that DNA-mediated fusion can be used to initiate the cell-free production of a model soluble and model membrane protein.

**Figure 1.**
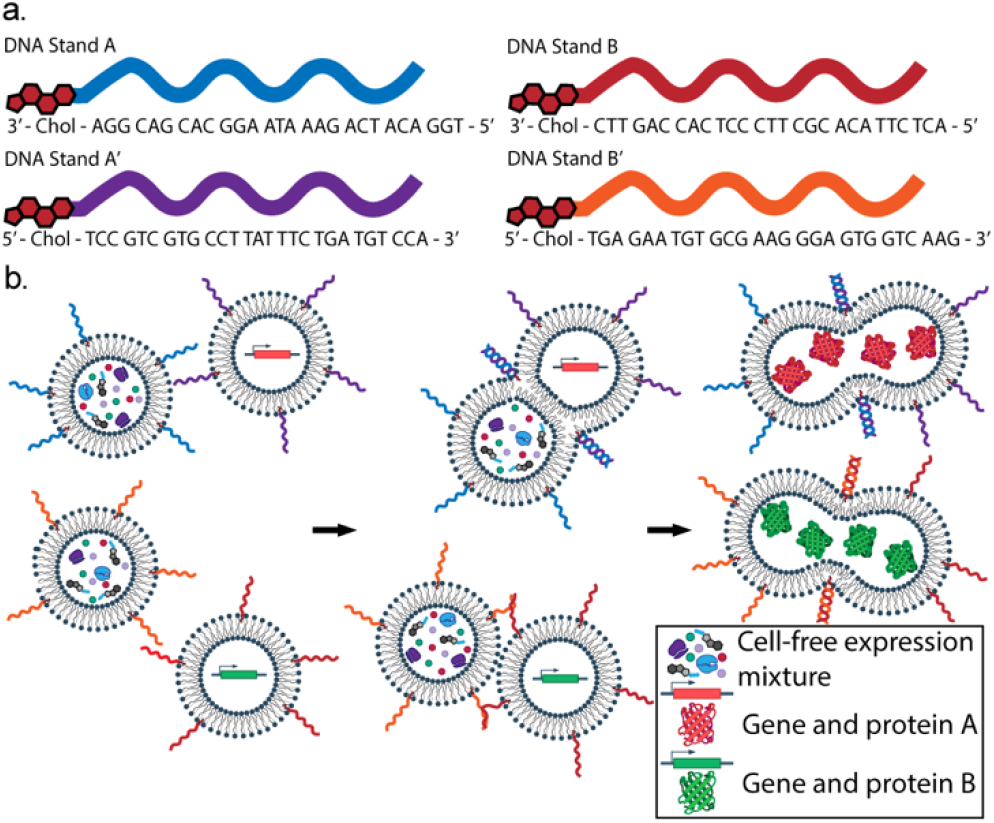
DNA oligonucleotides allow for controlled vesicle fusion. (a) Schematic of DNA oligonucleotide sequences used in this study. Two sets of complementary oligonucleotides (ex. A and A′) lead to vesicle fusion. (b) By utilizing different sets of oligonucleotides, vesicle fusion between specific vesicles may be controlled and allow for parallel biological reactions, such as cell-free protein synthesis, to occur.

## Results

### Membrane domains promote DNA-mediated membrane fusion

To assemble vesicles capable of fusion, we prepared small unilamellar lipid vesicles (SUVs) modified with DNA-lipid conjugates. The base composition of vesicle membranes was a homogenous, ternary mixture of dioleoyl-phosphatidylcholine (DOPC), dioleoyl-sn-glycero-3-phosphoethanolamine (DOPE), and cholesterol (Chol) at a molar ratio of 3:1:1. This composition was chosen as DOPE has been shown to aid fusion by introducing negative curvature to bilayer membranes,^[29,30]^ while cholesterol alters membrane organization and physical properties,^[31,32]^ and has been shown to aid fusion.^[33–38]^ After formation, vesicles were functionalized with cholesterol-conjugated oligonucleotides based on the design of Stengel and coworkers^[8,14]^ that enables complementary oligonucleotides (27mer) to hybridize in a zipper-like fashion, bringing opposing vesicles into contact (Figure 1 A, Table S1). To assess vesicle fusion, we utilized a total lipid-mixing assay that measures Förster resonance energy transfer (FRET) between two fluorescently-labelled phospholipids. Vesicles containing FRET-labelled lipids report membrane fusion between fluorescently labelled and unlabelled vesicles through a change in FRET efficiency that accompanies membrane growth.^[39,40]^ Using a standard curve relating mol % dye to FRET efficiency (Figure S1), we reported the percent lipid mixing as the change in mol % dye in vesicle systems. Using this assay, we first verified that vesicles modified with complimentary DNA strands were fusogenic (Figure 2). Reporter vesicles, containing FRET dyes, were mixed at a 1:4 molar ratio with target vesicles, lacking dyes, or an equivalent volume of PBS. We estimated that reporter vesicles were functionalized with ~100 DNA-oligonucleotides per vesicle, while target vesicles were functionalized with ~50 DNA-oligonucleotides per vesicle (see S.I. for calculation). We observed increases in FRET signal when vesicles contained complimentary DNA, but no changes in FRET signal when vesicles contained noncomplimentary (nc) DNA or no DNA tethers, confirming the necessity of complementary DNA tethers for inducing vesicle fusion.

**Figure 2.**
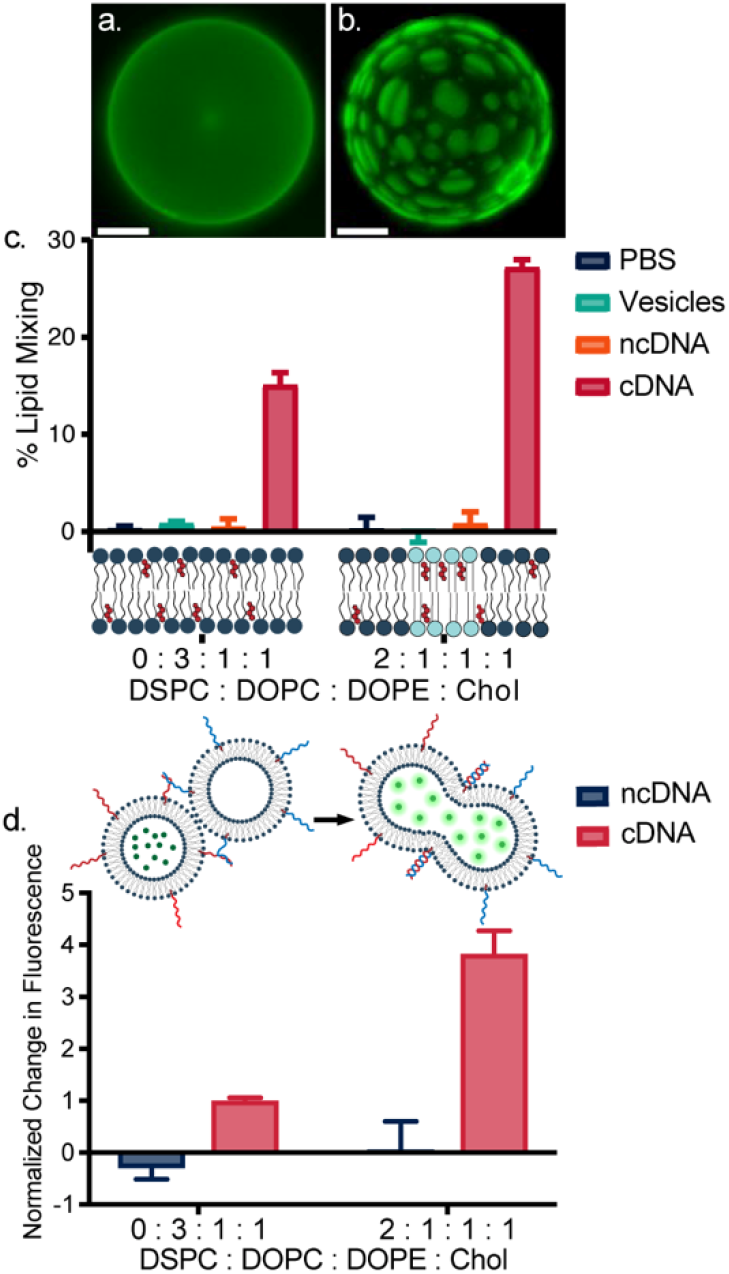
Phase segregation enhances DNA-mediated fusion. Maximum intensity projection of z-stacked fluorescence micrographs of GUVs containing DSPC: DOPC: DOPE: Chol at a molar ratio of (a) 0:3:1:1 and (b) 2:1:1:1 and with 0.1 mol% Rhod-PE, which localizes to the liquid-disordered phase. Scale bars are 5 μm. (c) Total lipid mixing for homogenous (0 DSPC: 3 DOPC: 1 DOPE: 1 Chol) and phase segregated vesicles (2 DSPC: 1 DOPC: 1 DOPE: 1 Chol). cDNA total lipid mixing for both lipid mixtures was significantly different from respective controls and from one another (p < 0.0001, Two-Way ANOVA Tukey Test). (d) Content mixing assay for homogenous and phase segregated vesicles reports increases in calcein fluorescence that accompany vesicle fusion. Values are normalized to the percent increase in calcein fluorescence observed upon mixing of homogenous vesicles functionalized with cDNA tethers. ncDNA and cDNA were not significantly different for 0 DSPC: 3 DOPC: 1 DOPE : 1 Chol (p > 0.05, Sidak’s Two-Way ANOVA), cDNA of 2 DSPC: 1 DOPC: 1 DOPE: 1 Chol was significantly different than all other samples (Sidak’s Two-Way ANOVA). Lipid and content mixing assays were carried out at 25°C at pH 7.3 in PBS. Error bars represent standard error of the mean (S.E.M), *n*=3.

We next explored the extent to which membrane domains could enhance membrane fusion between phospholipid vesicles. Recent studies with model membranes and computational studies have shown that the presence of phase-segregated domains in lipid vesicles could enhance fusion induced by an HIV fusion peptide.^[21,25,27,41]^ These studies demonstrated that membrane phase heterogeneity and line tension are important for viral membrane fusion. We wondered if membrane domains could be similarly used to enhance DNA-mediated vesicle fusion. In order to generate phase-segregated membranes, we incorporated distearoyl-phosphatidylcholine (DSPC) into lipid vesicles at a molar ratio of 2 DSPC:1 DOPC:1 DOPE:1 Chol. The inclusion of DSPC, which has 18 carbon acyl chains like DOPC and DOPE but is fully saturated, leads to lipid phase separation between liquid-ordered and liquid-disordered phases in the membrane.^[42]^ We confirmed the formation of membrane domains upon the inclusion of DSPC lipids through microscopy of giant unilamellar vesicles (GUVs) (Figure 2 A, B) and a bulk FRET assay for SUVs (Figure S2). Next, we used the total lipid mixing assay to monitor the effect of membrane domains on vesicle fusion. When vesicle membranes contained both domain-forming DSPC lipids and cDNA tethers, total lipid mixing during fusion increased from 15% to 27% (Figure 2 C). Interestingly, this enhancement in lipid mixing was only observed when both target and reporter vesicle pairs had lipid rafts (Figure S3). This result suggests that the increase in the extent of fusion is likely due to the thermodynamic instability of the presence of a two-phase membrane. It has been previously shown that liquid-ordered phases are generally thicker than the liquid disordered phase,^[22]^ and the hydrophobic mismatch has an associated energetic cost.^[23]^ As vesicles fuse, domains on opposing membranes are also likely to fuse,^[25]^ thus reducing the effective length of phase boundaries and reducing the free energy of the system overall. We again did not observe lipid mixing when vesicles containing non-complimentary tethers were mixed, regardless of the presence of membrane domains. Our results using total lipid mixing assays show that the presence of liquid-ordered phases improves the efficiency of DNA-meditated vesicle fusion.

To confirm that the observed lipid mixing initiated by DNA-mediated vesicle fusion corresponded to content mixing of encapsulated components, we performed a content calcein mixing assay. Vesicles with homogenous or phase segregated membranes were prepared encapsulating calcein at self-quenching concentrations and were mixed with empty vesicles of a similar membrane composition. Fusion and subsequent content mixing led to calcein dilution and de-quenching that could be monitored by measuring increases in calcein fluorescence. We observed that content mixing increased in vesicles that had phase segregated domains (2 DSPC: 1 DOPC: 1 DOPE: 1 Chol) relative to vesicles with homogenous membranes (3 DOPC: 1 DOPE: 1 Chol), indicating that the extent of total lipid mixing correlated with the extent of content mixing (Figure 2 D).

We then examined if hydrophobic mismatch between membrane domains, which should increase the line tension of membrane domains,^[23]^ could be used as a strategy to further enhance membrane fusion. We systematically varied the lipid acyl chain length of one membrane phase while maintaining the lipid acyl chain length in the other phase (18 carbon) and observed the impact of hydrophobic mismatch on total lipid mixing. First, we altered the thickness of the liquid ordered domain by replacing DSPC (18:0) with the shorter DMPC (14:0) and longer 24:0 PC lipid (Figure 3 A). Next, we varied the height of the liquid disordered domains by replacing DOPC (18:1) with the shorter 14:1 PC and longer 24:1 PC lipids (Figure 3 B). In either case, we observed that total lipid mixing increased as the lipid acyl chain length increased from 14 to 24 carbons. We observed via microscopy and FRET studies (Figure S2) that the longer 24 hydrocarbon chains generated larger domains with respect to lipids with 14 or 18 hydrocarbon chains. This phenomenon relating domain size to lipid length has been previously observed in model membranes.^[25,43]^ Increases in domain size are likely due to an increase in line tension, which arises from the difference in membrane thickness between the liquid-ordered and liquid-disordered phases.^[44,45]^ The total lipid mixing and FRET data indicate that increasing both the size of membrane domains and hydrophobic mismatch between domains increases the propensity of vesicles to fuse (Figure 3) and is consistent with previous work exploring fusion induced by HIV fusion peptides.^[25]^ Additionally, longer lipid tails may fill intermembrane voids that are created during membrane fusion, allowing the membrane to more easily bend and reform, and decreasing the energy barrier to achieve fusion.^[29,46]^ Notably, when domain formation was prevented by using only unsaturated lipids but hydrophobic mismatch was retained, total lipid mixing efficiency decreased to levels observed in homogenous vesicles with uniform membrane thickness (Figure 3 C). Taken together, these studies reveal that increasing the size of phase segregated domains and increasing hydrophobic mismatch between domain boundaries is a viable strategy to significantly enhance DNA-mediated vesicle fusion.

**Figure 3.**
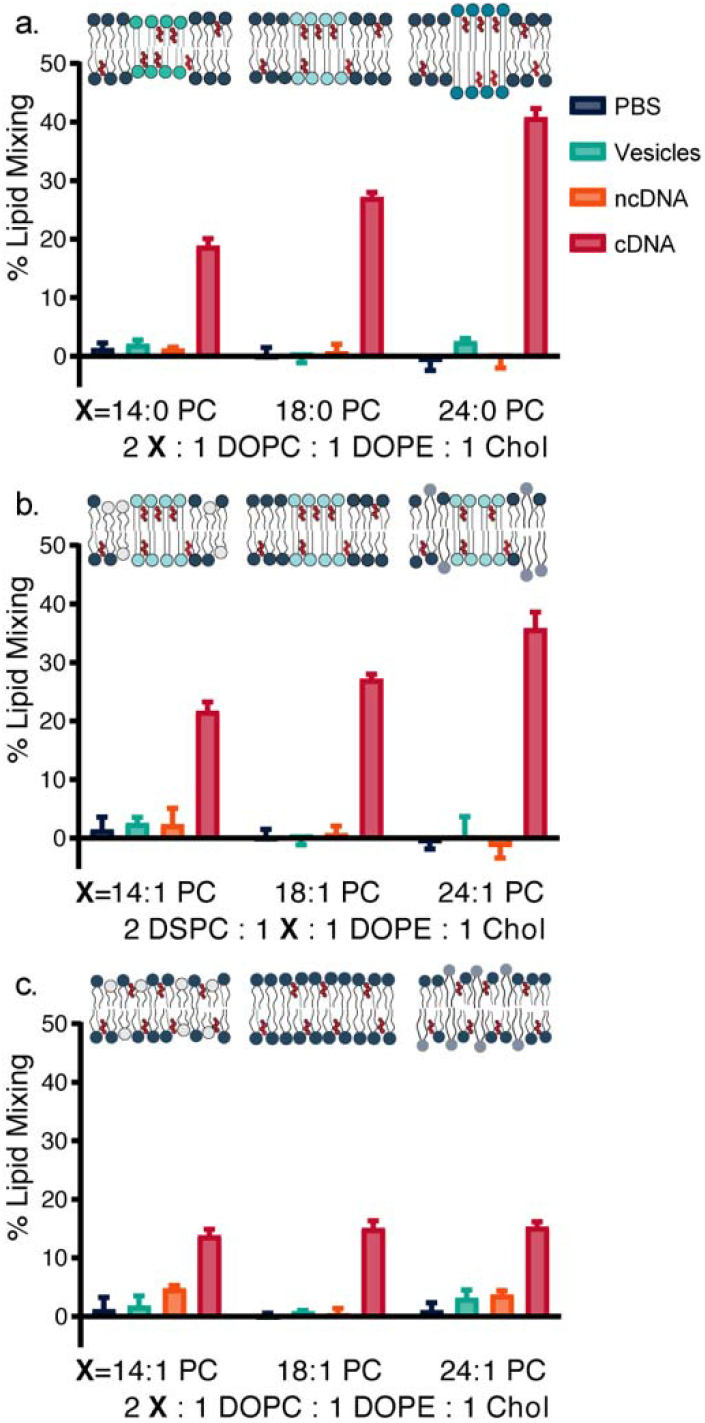
Hydrophobic mismatch controls DNA-mediated fusion in phase-segregated vesicles. (a) Total lipid mixing of vesicles in which the liquid ordered phase (X) is altered. Vesicles were composed of a molar ratio of 2:1:1:1 14:0 PC/DOPC/DOPE/Chol, 18:0 PC/DOPC/DOPE/Chol, and 24:0 PC/DOPC/DOPE/Chol. cDNA total lipid mixing for all lipid mixtures was significantly different from respective controls and from one another (p < 0.0001). (b) Total lipid mixing of vesicles in which the liquid disordered phase (X) is altered. Vesicles were composed of a molar ratio of 2:1:1:1 DSPC/14:1 PC/DOPE/Chol, DSPC/18:1 PC/DOPE/Chol, and DSPC/24:1 PC/DOPE/Chol. cDNA total lipid mixing for all lipid mixtures was significantly different from respective controls (p < 0.0001). 14:1 and 18:1 PC cDNA total lipid mixing were not significantly different from one another (p = 0.11), while 24:1 PC total lipid mixing was significantly different from all other cDNA lipid mixing samples. (c) Total lipid mixing of homogenous vesicles composed of liquid disordered lipids at a molar ratio of 2:1:1:1 14:1 PC/DOPC/DOPE/Chol, 18:1 PC/DOPC/DOPE/Chol, 24:1 PC/DOPC/DOPE/Chol. cDNA total lipid mixing for all lipid mixtures was significantly different from respective controls (p < 0.0001), but not from each other, as expected. Lipid mixing assays were carried out at 25°C at pH 7.3 in PBS. A two-way ANOVA followed by Tukey’s post hoc test was used. Error bars are S.E.M. for 3 replicates.

### DNA-mediated vesicle fusion initiates cell-free synthesis of proteins

We wondered if DNA-mediated fusion and the associated mixing of vesicle contents could be used to initiate a biological reaction in specific populations of vesicles. The use of liposomal compartments to engineer genetically-encoded reactions has been a long-standing goal in the design of artificial cellular systems. Recent work demonstrated how liposome fusion can mediate biological reactions.^[5,6]^ These studies utilized SNARE proteins and charge-charge interactions to initiate vesicle fusion, respectively, that provide an efficient method to drive vesicle fusion, but currently lack orthogonality that would allow for specific vesicle populations to be mixed simultaneously. Towards achieving this latter goal, we set out to determine the capacity of DNA-tethered vesicles to initiate protein synthesis in select populations of vesicles by inducing vesicle fusion.

We first investigated the extent to which DNA-tethers facilitated targeted fusion and cargo delivery to specific vesicle populations, features that would be important for initiating and sustaining biochemical reactions in vesicles. First, we used a lipid mixing assay to demonstrate we could direct when and where four distinct populations of vesicles fused with target vesicles present in the same solution. Target SUVs composed of phospholipids at a molar ratio of 3 DOPC: 1 DOPE : 1 Chol were prepared and functionalized with two different oligos (A and B) on their surface. Four populations of SUVs with a composition of 3 DOPC : 1 DOPE : 1 Chol were then added to this target population at two distinct time points. By varying the identity of the oligo on the added vesicles and the time at which vesicles were added, we were able to temporally control when and where fusion occurred, executing two stages of fusion in one vesicle population and one stage or no fusion with the others (Figure 4 A). The specificity of DNA-oligonucleotide binding was also supported by free energy calculations of DNA hybridization (Table S2). Towards the goal of initiating protein synthesis in vesicles, we then directly visualized the capacity of our system to deliver DNA to target vesicles using microscopy. We produced giant unilamellar vesicles (GUVs), composed of 2 DSPC : 1 DOPC: 1 DOPE : 1 Chol GUVs labelled with either rhodamine or Cy5.5 membrane dyes, that served as the target vesicles to which cargo would be delivered. SUVs were then prepared encapsulating a fluorescently-labelled 10-mer oligonucleotide (FAM-dA_10_) that represented genetic cargo that would need to be delivered to vesicles to initiate protein expression. We observed that SUVs fuse and deliver FAM-dA_10_ only to vesicles surface-modified with complimentary oligos, and do not fuse and deliver cargo to vesicles that have non-complimentary oligos on their surface. (Figure 4 B, C, S4, S5, S6). Together, these studies demonstrate the capacity for DNA-functionalized vesicles to mediate the temporally and spatially controlled delivery of vesicle cargo to specific vesicle populations.

**Figure 4.**
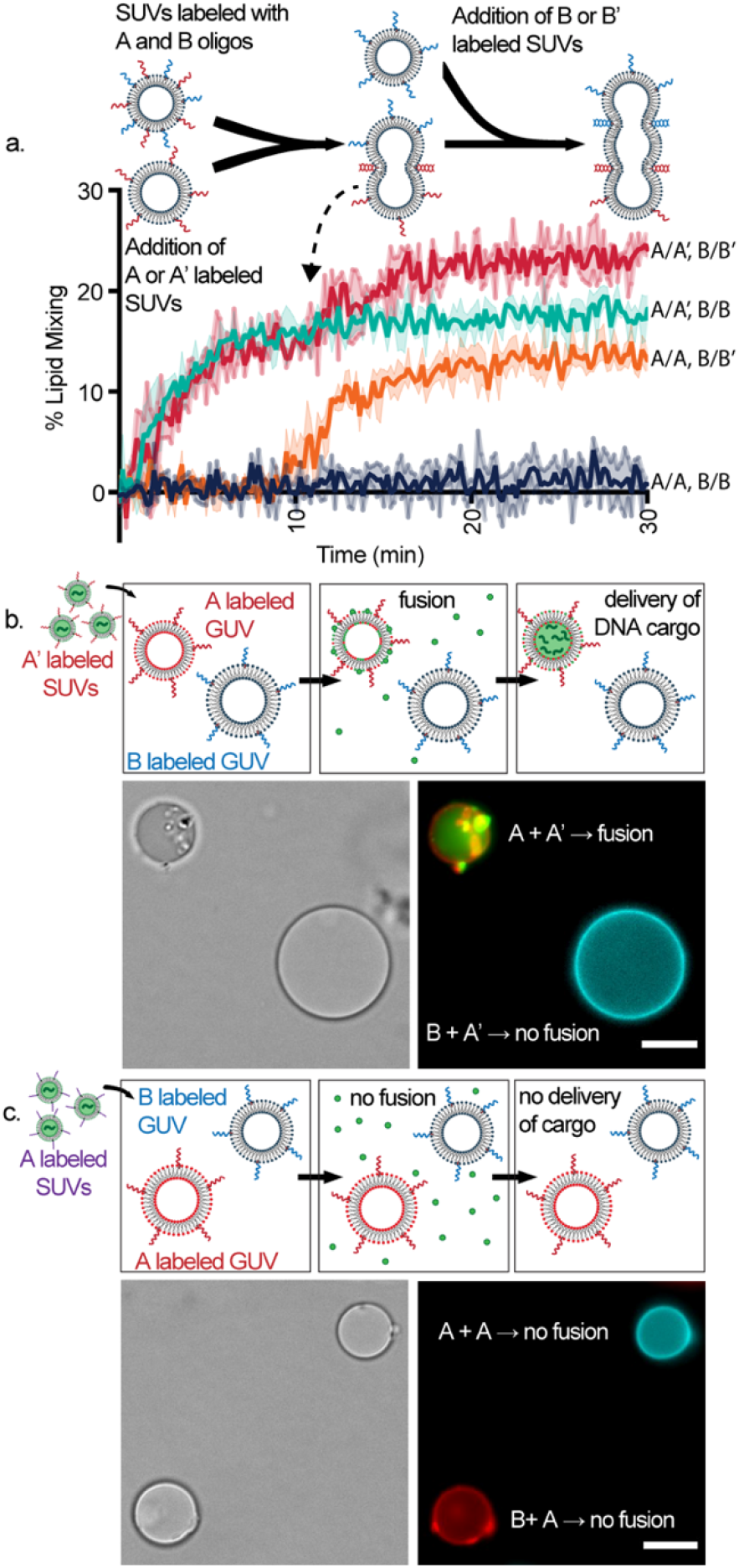
DNA-mediated fusion allows for the controlled fusion of specific vesicle populations and the delivery of DNA cargo. (a) A FRET total lipid mixing assay demonstrates that DNA oligonucleotides are orthogonal. SUVs composed of 3:1:1 DOPC/DOPE/Chol and labeled with Cy5.5/Cy7 membrane dyes were functionalized with ~100 strands of oligo A and B. At time t = 0 minutes, vesicles with oligo A or A′ were added, and at time t=10 minutes, vesicles with oligo B or B′ were added. This sequence of addition led to vesicles that underwent two fusion events (A/A′ and B/B′, red), one initial fusion event (A/A′ and B/B, light blue), a delayed fusion event (A/A and B/B′, orange), and no fusion events (A/A and B/B, dark blue). Lower and upper bounded lines represent the S.E.M., n=3. (b, c) DNA mediated fusion allows for the delivery of DNA-oligonucleotide cargo in specific populations of vesicles. Rhodamine labeled GUVs (red) with oligo A and Cy5.5 labeled GUVs (blue) functionalized with oligo B were mixed with SUVs encapsulating a fluorescently labeled FAM-dA_10_ oligonucleotide (green) and NBD PE membrane dye (green). When SUVs were functionalized with the complementary A′ oligo, they fused and delivered DNA cargo to the rhodamine GUVs (b). When SUVs were functionalized with the same oligo (a) as the rhodamine labeled GUVs, no fusion was observed (c). Representative DIC and fluorescent images are shown for each study. Microscopy studies were carried out at 25°C in PBS, pH 7.3. Scale bars are 10 μm.

We then investigated if content mixing via DNA-mediated vesicle fusion could initiate a complex biological reaction: cell-free protein synthesis. To maintain the stability of biological components used in the reaction, vesicles were prepared and maintained at 4°C, which is below the phase transition temperature of the phase segregated membranes, until the reaction time. Despite vesicle preparation being carried out at 4°C, domains formation and vesicle fusion appeared to be unaffected (Figure S7, S8) relative to vesicles that were prepared at 37°C. We first initiated a model enzymatic reaction by producing the fluorescent molecule resorufin. Briefly, we encapsulated the substrate, glucose, in one set of vesicles and glucose oxidase, horse radish peroxidase, and Amplex Red in another. Upon fusion, glucose was oxidized by glucose oxidase to yield hydrogen peroxide, which then oxidized Amplex Red in the presence of horseradish peroxidase to form the fluorescent molecule, resorufin (Figure S9). Next, we sought to synthesize model soluble and membrane proteins through the cell-free expression of superfolder green fluorescent protein (sfGFP) (Figure 5 A, S10) and a GFP fusion of the mechanosensitive channel of large conductance (MscL-GFP) (Figure 5 B), respectively. We encapsulated the plasmid encoding the protein of interest in one population of vesicles and a cell-free expression system (PURExpress) in another population of vesicles, and functionalized each set of vesicles with DNA tethers. Vesicles containing plasmid and PURExpress were brought to 37°C and mixed at a 1:1 ratio. We tracked protein synthesis by monitoring GFP fluorescence over time. We observed that vesicles that were surface-functionalized with cDNA tethers expressed protein products while those with ncDNA tethers did not. Furthermore, vesicles with phase segregated domains and cDNA tethers displayed significantly higher levels of fluorescence compared to non-phase segregated vesicles, demonstrating that increases in vesicle fusion efficiency that enhances content mixing has a direct effect on the yield of biochemical products. These results demonstrate that DNA-mediated fusion of synthetic vesicles may be used to initiate complex biological reactions in targeted vesicles. Furthermore, this study underscores how biophysical features of the membrane, which enhance membrane fusion, may be utilized to design and control the efficiency of biological reactions in vesicle systems.

**Figure 5.**
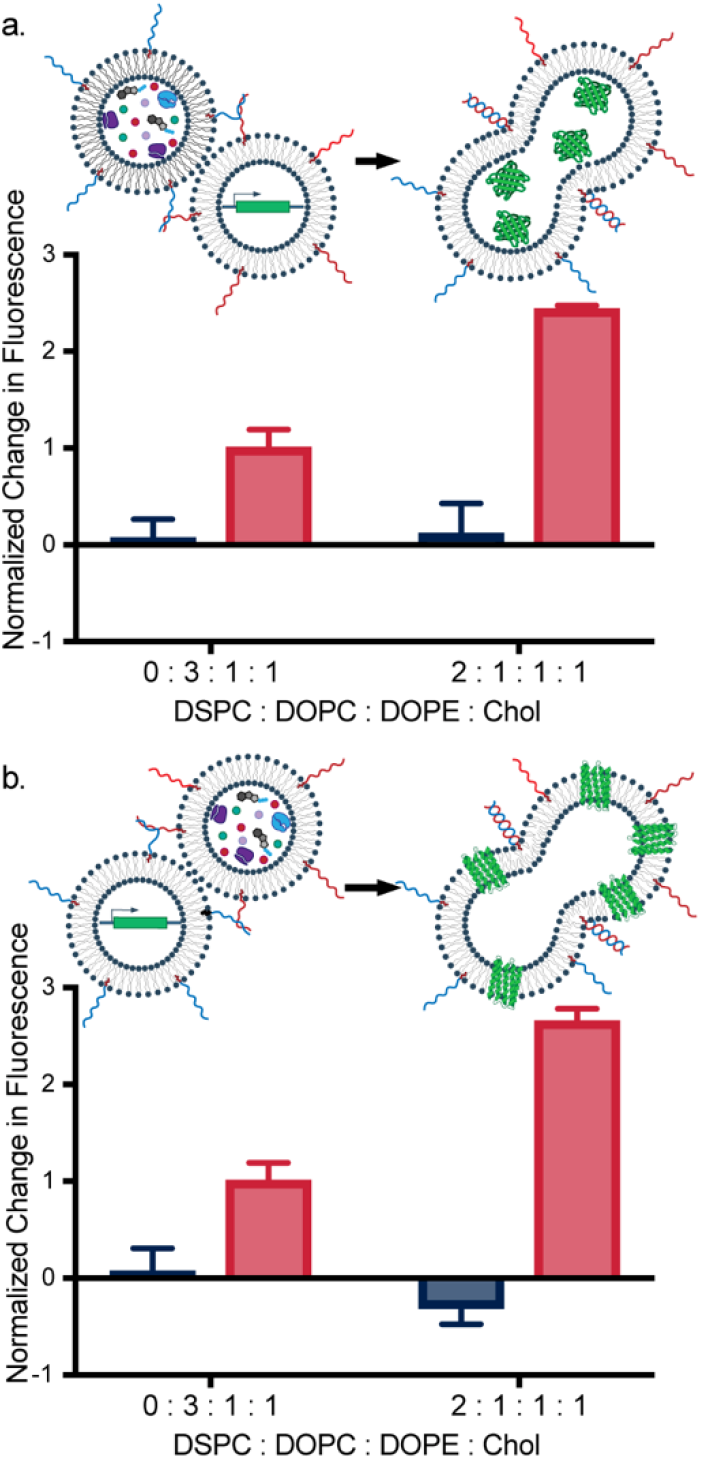
DNA-mediated vesicle fusion enables cell-free protein expression. Vesicle fusion and content mixing leads to the production of (a) a soluble protein, sfGFP, and (b) a membrane protein, MscL-GFP. ncDNA-functionalized vesicles (blue bars) do not fuse and thus protein fluorescence remains relatively unchanged, while vesicles functionalized with cDNA (red bars) fuse resulting in increased protein fluorescence as proteins are synthesized. Compositions of vesicle membranes that enhance fusion lead to increased protein production (a, b). Data is presented as the percent difference in fluorescence between the initial and final time point, normalized to the fusion of non-phase segregated vesicles. cDNA increase in fluorescence for all lipid mixtures was significantly different from respective controls and from one another (p < 0.02, Sidak’s Two-Way ANOVA). Error bars are S.E.M., *n=3*. Cell free expression studies were carried out at 37°C in HEPES buffer, pH 7.3.

## Discussion

Advances in biotechnology have enabled the break-down of biological systems into individual parts and the re-engineering and assembly of these parts, from the bottom-up, to design new materials. Here, lipid membranes and DNA tethers serve as fundamental biological parts that can be assembled in order to construct materials with new capabilities. By utilizing the molecular specificity of DNA hybridization, we were able to co-localize distinct populations of vesicles in an aqueous environment and drive their fusion and delivery of cargo. We were able to control the degree to which membrane fusion and content mixing occurred by assembling membranes of varying compositions. Finally, by leveraging the capacity of DNA to not only act as tethers but also encode genetic information, we initiated biochemical reactions within vesicles. Taken together, this work highlights the potential for DNA-mediated vesicle fusion to initiate the synthesis of biochemical products, with spatial and temporal control, in complex environments.

An important feature of our system was the use of membrane domains to enhance DNA-mediated vesicle fusion. Understanding how to harness biophysical properties of bilayer membranes to expand their functionality beyond containment alone is an important step in the design of artificial cellular systems. Cellular plasma membranes are heterogeneous and contain a variety of distinct regions with varying biophysical properties that we can draw from to design such systems. These physical features of the membrane can influence biochemical events by mediating fusion or increasing the local concentration of membrane proteins to aid targeting and adhesion.^[47,48]^ Notably, we are increasingly uncovering new mechanisms to recapitulate these features in synthetic vesicles^[49,50]^ that should lead to new opportunities to control systemic behaviors (ex. fusion, targeting, signalling) for which they are designed.^[51]^

DNA-oligonucleotides tethered to vesicle membranes allow for the engineering of precise, targeted fusion events between vesicles. The nearly infinite array of molecularly specific DNA hybridization pairs allows for DNA-mediated fusion systems to be orthogonal to surrounding systems, surpassing the currently known available pairs of protein tethers. Utilizing DNA as surface-localized molecules to ‘barcode’ specific vesicles for fusion could allow for many vesicle fusion architectures to be constructed. Potential fusion schemes should allow for multiple, orthogonal reactions to be performed in parallel or for cascaded reactions to be performed by fusing multiple populations of vesicles to one set of target vesicles.

Advances in the field of synthetic biology have brought us closer to the goal of designing artificial cells and cellular mimetic structures capable of complex biomanufacturing.^[7,52–56]^ Genetic circuitry can be harnessed in cell-free environments in order to produce a variety of biological molecules.^[57,58]^ However, the synthesis of biological molecules with the diversity, efficiency, and spatial and temporal precision of cellular systems remains challenging. This challenge may be addressed through thoughtful engineering of synthetic membranes combined with strategic design of membrane architecture to allow for efficient, targeted, and specific fusion between vesicle compartments. We expect that by combining membrane design with improved DNA linker chemistry,^[15,16]^ biochemical product yield could be further improved.

Biochemical products synthesized within vesicles as a result of fusion could further expand the functionality of vesicles by generating useful protein products. These protein products could include pore-forming proteins to release and deliver cargo,^[6,59,60]^ or surface antigens that enable sensing of and interactions with the surrounding environment. Ultimately, DNA-mediated vesicle fusion coupled with the cell-free synthesis of biological molecules should expand the possible functions of vesicle-based materials, enabling enhanced sensing, delivery, and biomanufacturing capabilities.

## Conclusion

In summary, we demonstrated that DNA-mediated membrane fusion efficiency may be enhanced by altering vesicle membrane composition to include membrane domains. By altering hydrophobic thickness within each membrane phase, we were able to further tune fusion efficiency. Harnessing the specificity and orthogonality of DNA oligonucleotide hybridization, we then showed DNA-mediated vesicle fusion can be used to deliver DNA-based cargo to different vesicle populations. Lastly, we demonstrated that DNA-mediated vesicle fusion allows for the delivery of biochemical reactants and initiation of model biological reactions by initiating an enzymatic reaction and the *in vitro* production of a model soluble and membrane protein. Taken together, this work demonstrates how DNA-mediated vesicle fusion may allow for the engineering of encapsulation systems which support the initiation and spatial control of complex reactions. Continuing to develop such technologies will allow for the design and development of more advanced cell-free and artificial cellular systems, capable of performing complex tasks in varying environments ranging from the body to bioreactors.

## Experimental Section

### Materials

1,2-dioleoyl-sn-glycero-3-phosphocholine (DOPC), 1,2-dioleoyl-sn-glycero-3-phosphoethanolamine (DOPE), Cholesterol, 1,2-distearoyl-sn-glycero-3-phosphocholine (DSPC), 1,2-dimyristoyl-sn-glycero-3-phosphocholine (DMPC), 1,2-dilignoceroyl-sn-glycero-3-phosphocholine (24:0 PC), 1,2-dimyristoleoyl-sn-glycero-3-phosphocholine (14:1 PC), 1,2-dinervonoyl-sn-glycero-3-phosphocholine (24:1 PC), 1,2-dioleoyl-sn-glycero-3-phosphoethanolamine-N-(7-nitro-2-1,3-benzoxadiazol-4-yl)1,2-dioleoyl-sn-glycero-3-phosphoethanolamine-N-(lissamine rhodamine B sulfonyl) (18:1 Rhodamine), 1,2-dioleoyl-sn-glycero-3-phosphoethanolamine-N-(Cyanine 5.5) (Cy 5.5 PE), 1,2-dioleoyl-sn-glycero-3-phosphoethanolamine-N-(Cyanine 7) (Cy 7 PE) were purchased from Avanti Polar Lipids. 1,2-dipalmitoyl-sn-glycero-3-phosphoethanolamine-N-(7-nitro-2-1,3-benzoxadiazol-4-yl) was purchased from Invitrogen. Calcein dye, glass bottomed Lab-Tek II microscope chambers, 20K MWCO Slide-A-Lyzer Mini Dialysis Kits and the DNA-oligonucleotides were obtained from Thermo Fisher. Phosphate-buffered Saline (PBS), HEPES buffer components, sucrose, and Sepharose 4B (45−165 μm bead diameter) were obtained from Sigma Aldrich. FAM-dA10 was purchased from IDT (Coralville, IA). PURExpress was obtained from New England Biosciences. pJL1-sfGFP was a gift from Michael Jewett (Addgene plasmid no. 102634, Northwestern University, Evanston, IL),^[61]^ EcMscL gene (pADtet-His6-MscL) was a gift from Allen Liu (Addgene plasmid no. 83373, California Institute of Technology, Pasadena, CA),^[62]^ and mEGFP was a gift from Michael Davidson (Addgene plasmid no. 54622, Florida State University, Tallahassee, FL).

### Vesicle Preparation

Small unilamellar vesicles were prepared through a thin film hydration method.^[63]^ Lipid dissolved in chloroform was mixed in molar ratios specified in the manuscript. Chloroform was evaporated under a stream of nitrogen to create uniform lipids films, and films were placed under vacuum for 3 hours to remove any remaining chloroform. All vesicles were rehydrated overnight at 60°C, except for vesicles containing 24:0 PC, which were heated to 81°C (above its phase transition temperature of 80.3°C) or for vesicles prepared containing cell-free components, which were hydrated at 4°C. Vesicles for FRET and calcein assays were rehydrated in 290 mOsm PBS, pH 7.3. Vesicles for protein expression assays were rehydrated in PURE Buffer (50 mM HEPES, 100 mM KCl, 10 mM MgCl_2_). Vesicles were briefly vortexed and extruded using an Avanti miniextruder through a 0.1 μm polycarbonate filter for all FRET experiments and through a 0.4 μm polycarbonate filter for all content mixing and reaction experiments to seven passes. For FRET lipid mixing and calcein content mixing experiments, target vesicles were functionalized with 100 oligos per vesicle, while added vesicles were functionalized with 50 oligos per vesicle. For protein synthesis, both sets of vesicles were functionalized with 100 strands per vesicle. DNA-oligos were added to vesicles after hydration and processing and incubated at 25°C for 20 minutes before they were used in fusion studies.

### Total Lipid Mixing Assays

The extent of lipid mixing was determined by measuring the change in Forster resonance energy transfer (FRET) between two membrane dyes (Cy5.5/Cy7). SUVs were added to a cuvette in a molar ratio of 1:4 of labelled (1 mol% Cy5.5/7) to unlabeled vesicles to a final concentration of 200 μM. Fluorescence change was measured using an Agilent Cary Eclipse Fluorescence Spectrophotometer by exciting vesicle samples at 675 nm and recording the emission intensity at 713 nm (Cy5.5, F_donor_) and 775 nm (Cy7, F_acceptor_) at room temperature. Reporter vesicles containing FRET dyes were read for 2 minutes, which served as the “zero” time point, and read for a subsequent 30 minutes following the addition of target (unlabeled) vesicles. Vesicles were disrupted using Triton X-100 (1%) to measure the unquenched fluorescence of the donor. FRET measurements were normalized by FRET values obtained after vesicle lysis with 1% Triton-X.^[40]^

The FRET ratio was calculated as the following ratio:

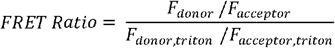

To quantify lipid mixing, a standard curve of vesicles containing 0, 0.15, 0.3, 0.5, 0.65, 0.8, 1.0 mol% dye was produced for each lipid mixture. An exponential fit was then performed on each standard curve in the form of y = e^Ax+B^, where A and B are constants, x is the FRET ratio, and y is the calculated mol% dye. Total lipid mixing values were then calculated using the fitted equation to calculate the effective mol% dye over time. The calculated mol% dye was then converted to total lipid mixing by using the following equation:

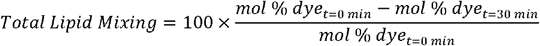

### Content Mixing

For content mixing assays, donor vesicles were rehydrated with 20 mM calcein in PBS. Calcein vesicles were purified using a size exclusion column packed with Sepharose 4B immediately before experimentation. Calcein vesicle fluorescence (ex. 480/em. 520 nm) was read for 2 minutes on a fluorescence spectrophotometer before adding empty vesicles in a 1:4 molar ratio of calcein to empty vesicles. Fluorescence of calcein was recorded for 30 minutes at room temperature. 1% Triton-X was then added to achieve a maximum dequenching of calcein, which served as the fluorescence intensity for 100% mixing. Percent content mixing was calculated using the following equation:

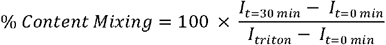

Where I_t=0 min_ is the initial fluorescence intensity, I_t=30 min_ is the fluorescence intensity at 30 minutes, and I_triton_ is the fluorescence intensity after the addition of Triton-X.

### Preparation of GUVs

GUVs were prepared via electroformation using the Nanion Vesicle Prep Pro (Nanion Technologies) standard vesicle preparation protocol. 10 mM mixtures of lipid in chloroform were prepared with 0.1 mol% Rhod-PE. 10 μL of each solution was then drop-casted onto indium tin oxide slides and placed under vacuum for 20 minutes to eliminate solvent. Films were rehydrated with 290 mOsm sucrose.

### Microscopy

GUVs were observed under a microscope to visualize domain formation. Glass bottomed Lab-Tek II microscope chambers (Thermo Fischer) were used to image GUVs. 200 μL of bovine serum albumin was placed into each chamber and allowed to sit for 30 min. Each well was then washed with 290 mOsm PBS and 1 μL of 1 mM of GUVs were added to 250 μL of PBS and allowed to settle in each chamber. A 40x Oil objective was used to visualize vesicles. Images were acquired on a Nikon Eclipse Ti2 Inverted Microscope and maximum intensity projection of Z-stacks were produced using NIS software to visualize domains.

### Visualizing DNA delivery to GUVs

To visualize vesicle fusion and delivery of FAM-dA_10_, GUVs composed of 1 DSPC: 1 DOPC: 1 DOPE: 1 Chol with either 0.1 mol% Rhod-PE or Cy5.5 PE were made via electroformation using the method outlined above. SUVs were composed of 1 DSPC: 1 DOPC: 1 DOPE : 1 Chol with 0.1 mol% NBD-PE and encapsulated 30 μM FAM-A_10_ and were extruded 7x to 400 nm. GUVs were functionalized with 10 μM of DNA-oligos while SUVs were functionalized with ~100 oligos per vesicle. GUVs and SUVs were mixed in a 1 to 10 molar ratio (1 mM final lipid concentration) for 1 hour. The vesicle solution was then diluted to 10 μM and 200 uL of the vesicle solution was then deposited into a Lab-Tek microscope chamber and imaged using a 40x oil objective using a Nikon Eclipse Ti2 Inverted Microscope.

### Cell Free Protein Expression

Lipid films were either rehydrated with 40 ng/μl of plasmid in PURE buffer (50 mM HEPES, 100 mM KCl, 10 mM MgCl_2_) or the PURExpress System and allowed to rehydrate and form vesicles for 4 hours at 4°C. Vesicles were then briefly vortexed and extruded 7x through a 400 nm filter. Both sets of vesicles were dialyzed against PURE buffer with 2 buffer exchanges overnight in a 20K MWCO Slide-a-lyzer cassette at 4°C. Each set of vesicles were then functionalized with 100 oligos per vesicle and incubated for 20 minutes. Vesicles were mixed in a 1:1 ratio, for a final concentration of 500 μM. Fluorescence for GFP (ex. 480/ em. 507 nm) was read for 4 hours at 37°C on a fluorescence spectrophotometer.

## Acknowledgements

We thank members of the Kamat Lab for thoughtful discussions. This research was supported in part by the Searle Funds at The Chicago Community Trust and the National Science Foundation under Grant No. 1844336. J.A.P. was supported by an NSF Graduate Research Fellowship. M.L.J. was supported by Grant Number T32GM008382 from the National Institute of General Medical Sciences.

