## Supporting Information for "Barcoding biological reactions with DNA-functionalized vesicles"

#### Table of Contents

Experimental Procedures ………….2
Supporting Tables and Figures……4
References…………………………..12

**Experimental Procedures**

**Materials.** 1,2-dioleoyl-sn-glycero-3-phosphocholine (DOPC), 1,2-dioleoyl-sn-glycero-3-phosphoethanolamine (DOPE), Cholesterol, 1,2-distearoyl-sn-glycero-3-phosphocholine (DSPC), 1,2-dimyristoyl-sn-glycero-3-phosphocholine (DMPC), 1,2-dilignoceroyl-sn-glycero-3-phosphocholine (24:0 PC), 1,2-dimyristoleoyl-sn-glycero-3-phosphocholine (14:1 PC), 1,2-dinervonoyl-sn-glycero-3-phosphocholine (24:1 PC), 1,2-dioleoyl-sn-glycero-3-phosphoethanolamine-N-(7-nitro-2-1,3-benzoxadiazol-4-yl)1,2-dioleoyl-sn-glycero-3-phosphoethanolamine-N-(lissamine rhodamine B sulfonyl) (18:1 Rhodamine) were purchased from Avanti Polar Lipids. 1,2-dipalmitoyl-sn-glycero-3-phosphoethanolamine-N-(7-nitro-2-1,3-benzoxadiazol-4-yl) and Amplex Red Glucose/Glucose Oxidase Kit were purchased from Invitrogen. The DNA-oligonucleotides were obtained from Thermo Fisher. PURExpress was obtained from New England Biosciences.

**Measuring domain size via FRET.** Lipid domain size in SUVs was indirectly measured via Förster resonance energy transfer (FRET). Using dyes which are conjugated with saturated and unsaturated lipid (18:1 Rhodamine (acceptor) and 16:0 NBD (donor)), donor and acceptor dyes preferentially segregate into the Lo and Ld phase, respectively. Domain size affects the average separation between donor and acceptor dyes in a membrane, altering the FRET efficiency and allows for the detection of relative changes in domain size. Fluorescence intensity of 200 μM vesicles was read using an Agilent Cary Eclipse Fluorescence Spectrophotometer by exciting the sample at 463 nm and recording the intensity at 517 nm (NBD, donor) and 590 nm (Rhodamine, acceptor). Fluorescence was reread upon the addition of 1% Triton-X. FRET efficiency was reported as the FRET ratio, calculated as $F_{donor}/F_{acceptor}$.

**Initiating Enzymatic Reaction via DNA-mediated fusion.** Two sets of 2 DSPC:1 DOPC:1 DOPE : 1 Chol vesicles were produced at a 20 mM concentration of total lipids: One set encapsulated 200 μM glucose, while the other encapsulated glucose oxidase, horse radish peroxidase, and Amplex Red to the concentrations specified by the Amplex Red Glucose/Glucose Oxidase Kit (Invitrogen). Rehydration was conducted at 4ºC. Vesicles were extruded through a 400 nm polycarbonate filter to 7 passes and purified via gravity size exclusion chromatography using a Sepharose 4B packed column. Vesicles were then functionalized with DNA-oligos and incubated for 20 minutes. Glucose and enzyme filled vesicles were mixed in a 3:1 molar ratio and diluted to 200 μM and allowed to react for 1 hour at room temperature before performing flow cytometry.

**Measurement of vesicle fusion reactions by flow cytometry.** A BD LSRII (BD Biosciences) flow cytometer was used to measure GFP and Amplex Red production upon vesicle fusion. Vesicles were loaded and gated for size between 400 nm and 1,000 nm to reduce any autofluorescence from large vesicle clumps and 10,000 events were recorded. GFP was excited with 488 nm laser and detected with 530/30 bandpass filter, while Amplex Red was excited with 561 nm laser and detected with 610/20 bandpass filter. Data was analyzed using FlowJo, and statistical analysis was conducted using a chi-squared test.

**Calculating the amount of DNA-oligonucleotides to add to vesicles to achieve desired DNA functionalization.** The amount of DNA-oligonucleotides to add to a given vesicle stock in order to functionalize vesicles, of radius R, with the desired number oligonucleotides was calculated using the following formulas:

$$Surface\ Area = 4\pi R^2$$

$$Chains\ per\ vesicle = \frac{surface\ area}{area\ per\ lipid\ chain} \times\ 2\ leaflets\ per\ vesicle$$

$$Vesicles\ per\ film = \frac{moles\ of\ lipid \times N_A}{chains\ per\ vesicle}$$

$$Moles\ of\ DNA\ required = number\ of\ oligos\ per\ vesicle \times \frac{vesicles\ per\ film}{N_A}$$

where $N_A$ is Avogadro's number.

It was assumed that the average area per lipid is 0.6 $nm^2$.[1]
Curvature of the vesicles was assumed to be negligible so that the inner and outer leaflet have identical surface areas and number of lipid chains.

### Supporting Tables and Figures

**Table S1.** DNA-oligonucleotide sequences. Two sets of 27 nucleotide sequences were used in this study. Oligonucleotides were conjugated to cholesterol separated by a TEG spacer.

| Name | Sequence (5' to 3') |
|---|---|
| A | TGG ACA TCA GAA ATA AGG CAC GAC GGA [CholTEG] |
| A' | [CholTEG] TCC GTC GTG CCT TAT TTC TGA TGT CCA |
| B | CTT GAC CAC TCC CTT CGC ACA TTC TCA [CholTEG] |
| B' | [CholTEG] TGA GAA TGT GCG AAG GGA GTG GTC AAG |

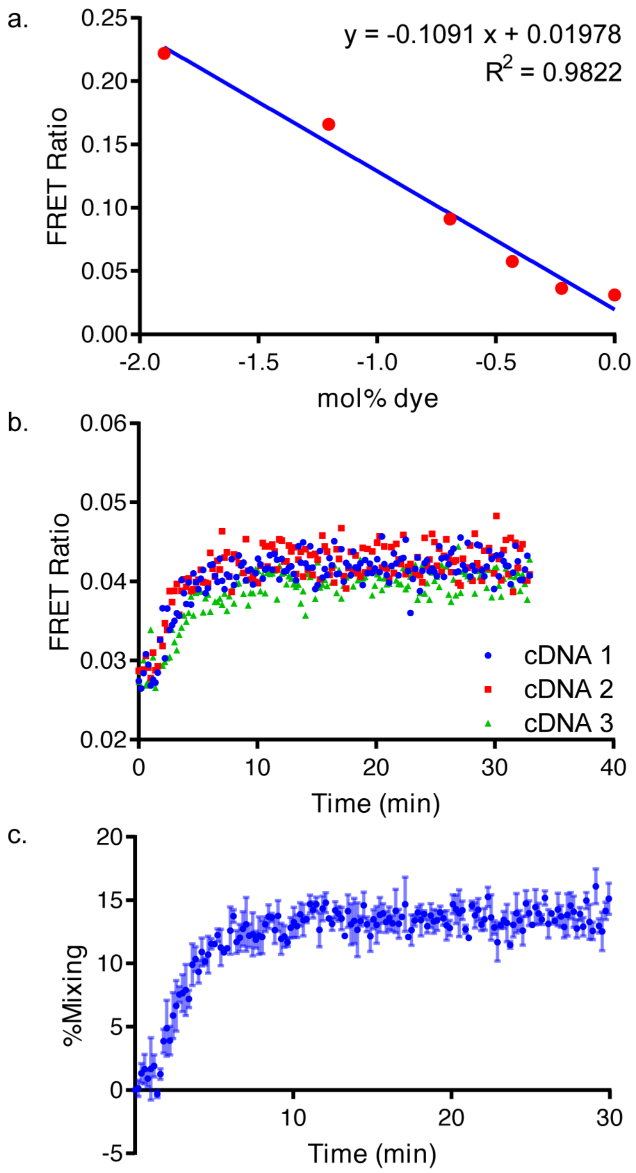


**Figure S1.** Data analysis for FRET total lipid mixing assay. (a) A standard curve was generated for each lipid composition. (b) FRET ratios ($F_{donor}/F_{acceptor}$) were then recorded for 30 minutes. (c) Using the standard curve, FRET ratios were converted into mol% dye. From this information, percent mixing was calculated.

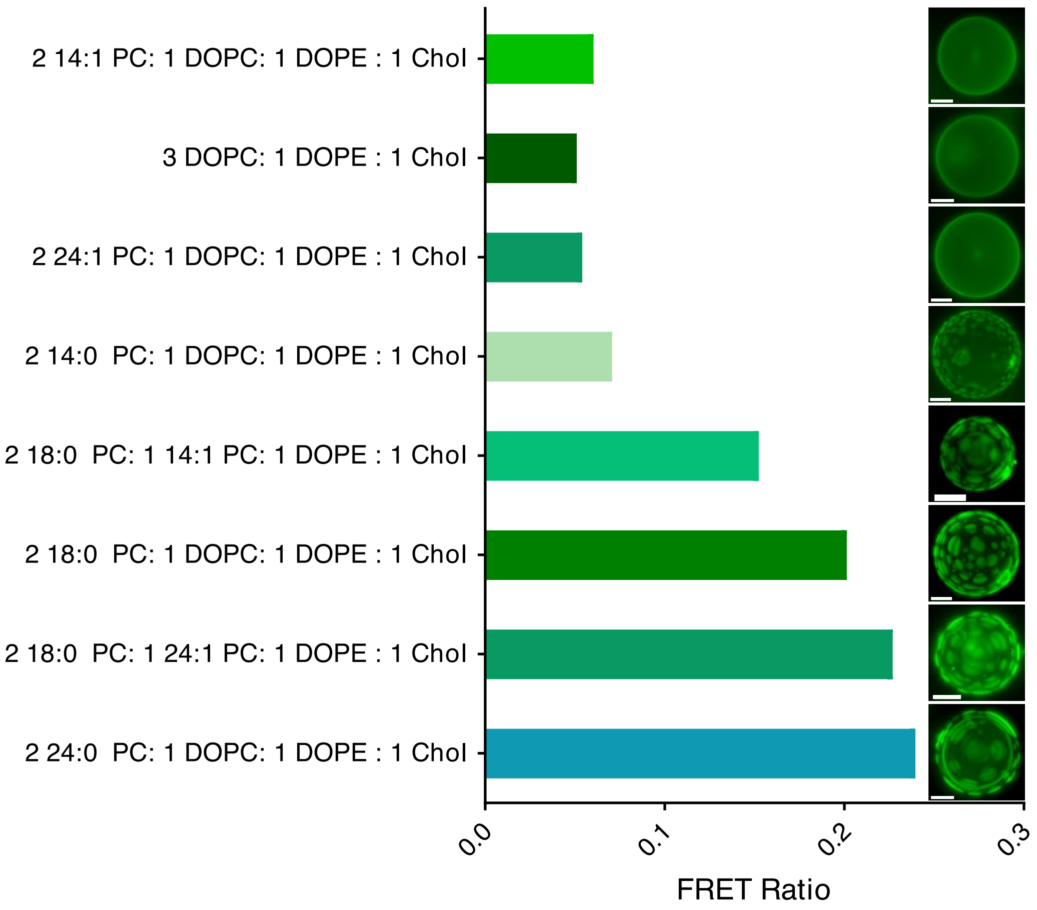


**Figure S2.** FRET Ratios and maximum intensity projection of z-stacked fluorescence micrographs of GUVs for all membrane mixtures studied. Bulk FRET ratios were measured using 18:1 Rhodamine and 16:0 NBD, expected to segregate into the liquid disordered and liquid ordered phase, respectively. Larger domains yielded larger FRET ratios. Z-stacked images of vesicles containing 0.1 mol% 18:1 rhodamine. Scale bars are 5 µm.

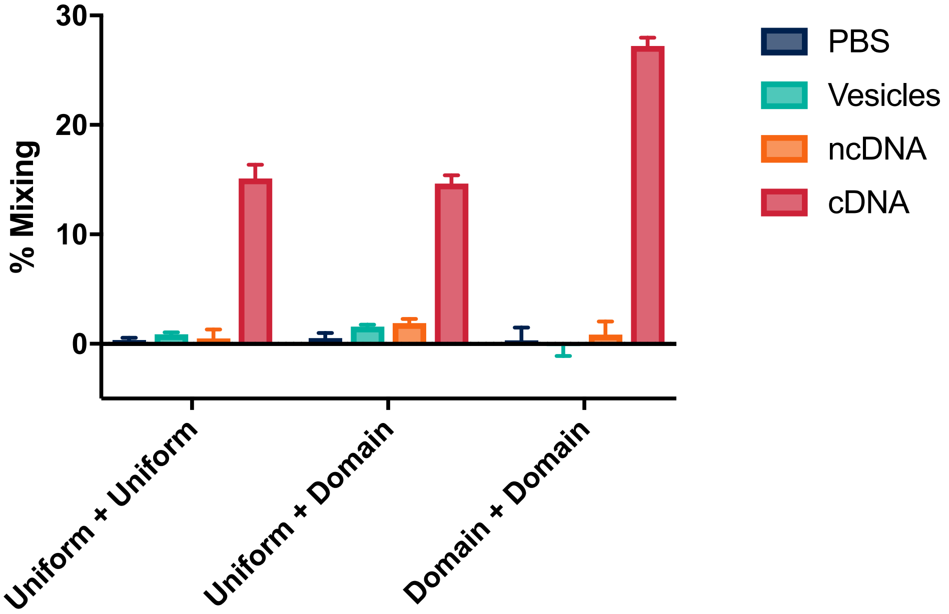


**Figure S3.** Fusion of vesicles with and without lipid domains. When one set of vesicles (uniform + domain) does not exhibit phase segregation, total lipid mixing is comparable to that observed with non-phase segregated vesicles containing complimentary DNA tethers (uniform + uniform). When both target and reporter vesicles containing cDNA tethers both exhibit phase-segregated domains (domain + domain), total lipid mixing nearly doubles. Uniform vesicles were composed of 3 DOPC :1 DOPE :1 Chol, while domain vesicles were composed of 2 DSPC : 1 DOPC : 1 DOPE : 1 Chol. Error bars represent S.E.M. for a sample size of 3.

**Table S2.** Gibbs Free energy of oligonucleotide binding. Complementary strands of DNA exhibit the most favorable conformation for oligo-oligo binding. Values are reported in kcal/mol.

| | *A* | *A'* | *B* | *B'* |
|---|---|---|---|---|
| ***A*** | -3.61 | **-52.23** | -6.61 | -6.44 |
| ***A'*** | **-52.23** | -3.61 | -6.44 | -6.61 |
| ***B*** | -6.61 | -6.64 | -3.61 | **-51.33** |
| ***B'*** | -6.64 | -6.61 | **-51.33** | -3.61 |

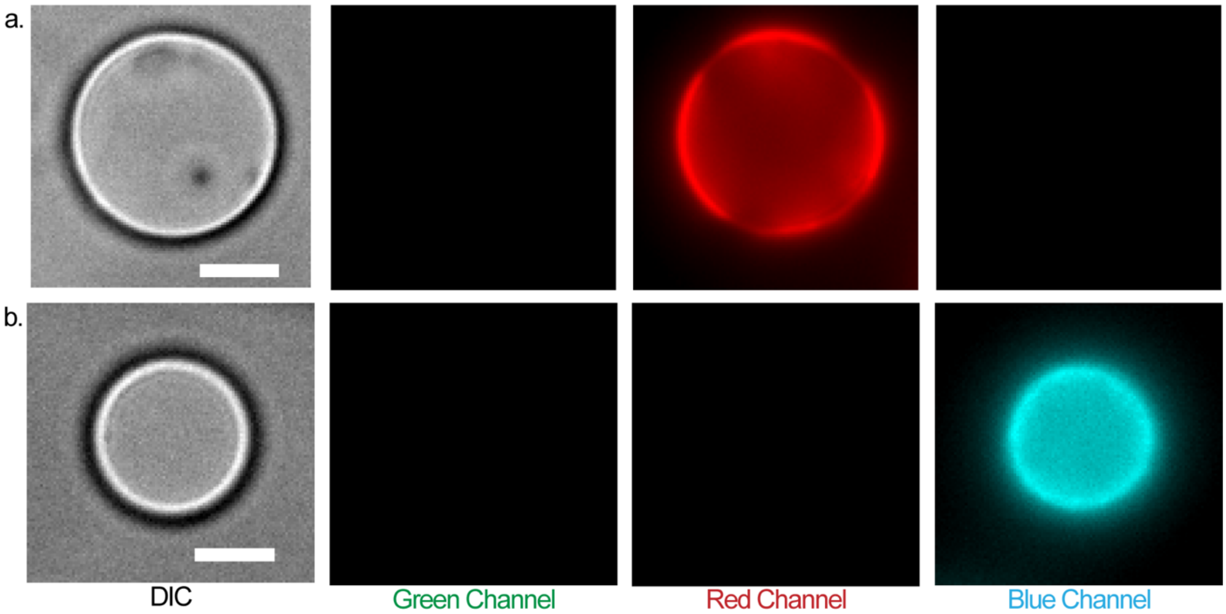


**Figure S4.** GUVs prior to the addition of SUVs do not fluoresce in the green channel. (a) Rhodamine labeled and (b) Cy5.5 labeled GUVs composed of 2:1:1:1 DSPC/DOPC/DOPE/Chol. Each column is a different fluorescent channel. Vesicles do not uniformly fluoresce as the lipid dyes are conjugated to an unsaturated lipid, and therefore segregate into the Ld phase. Scale bars are 5μm.

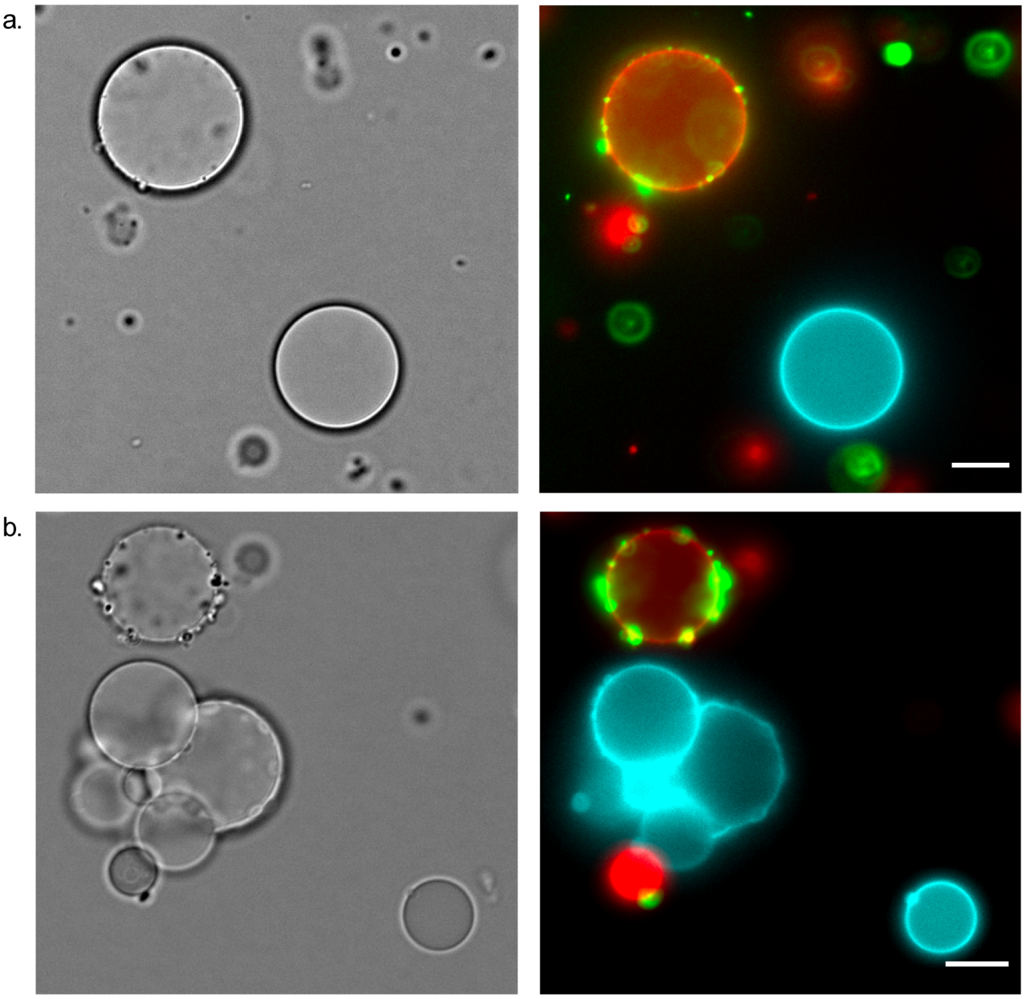

**Figure S5.** Hemifusion of SUVs was observed to GUVs, in addition to full fusion. DIC (left) and overlay of red, green, and blue fluorescent channels (right) of SUVs (labeled with A′) hemifused to rhodamine labeled GUVs (labeled with oligo A). SUVs only hemifused with rhodamine labelled GUVs as SUVs were functionalized with cDNA tethers to rhodamine labeled GUVs, but not Cy5.5 labeled GUVs (labeled with oligo B). Scale bars are 10 μm.

a.

b.

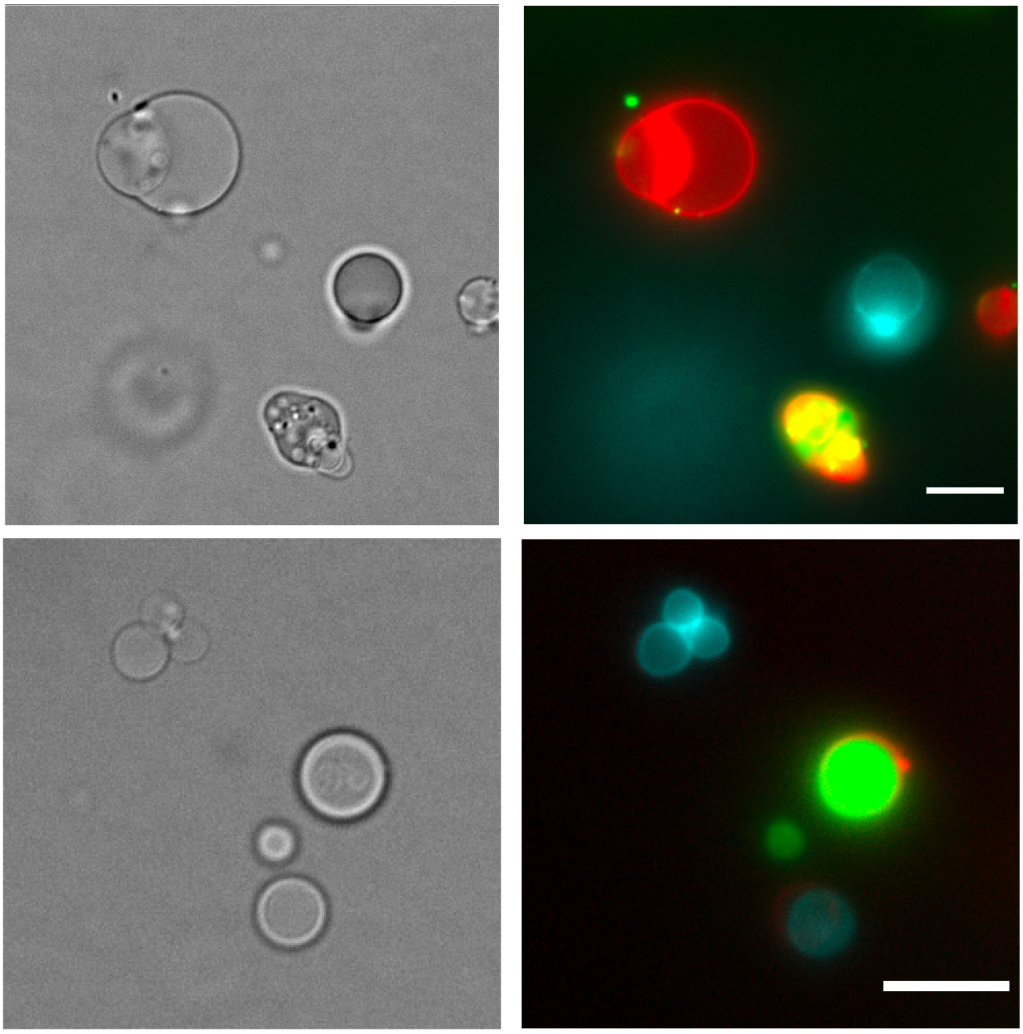

**Figure S6.** Delivery of FAM-$dA_{10}$ to GUVs. DIC (left) and overlay of red, green, and blue fluorescent channels (right) of SUVs which have fused and delivered FAM-$dA_{10}$ to rhodamine labeled GUVs, but not Cy5.5 labeled GUVs. Fusion was only observed between SUVs (labeled with A′) and rhodamine labelled GUVs (labeled with oligo A) as they were labelled with cDNA tethers. Cy5.5 labelled GUVs were not labelled with cDNA (labeled with oligo B) tethers and therefore no fusion and delivery was observed. Scale bars are 10 μm.

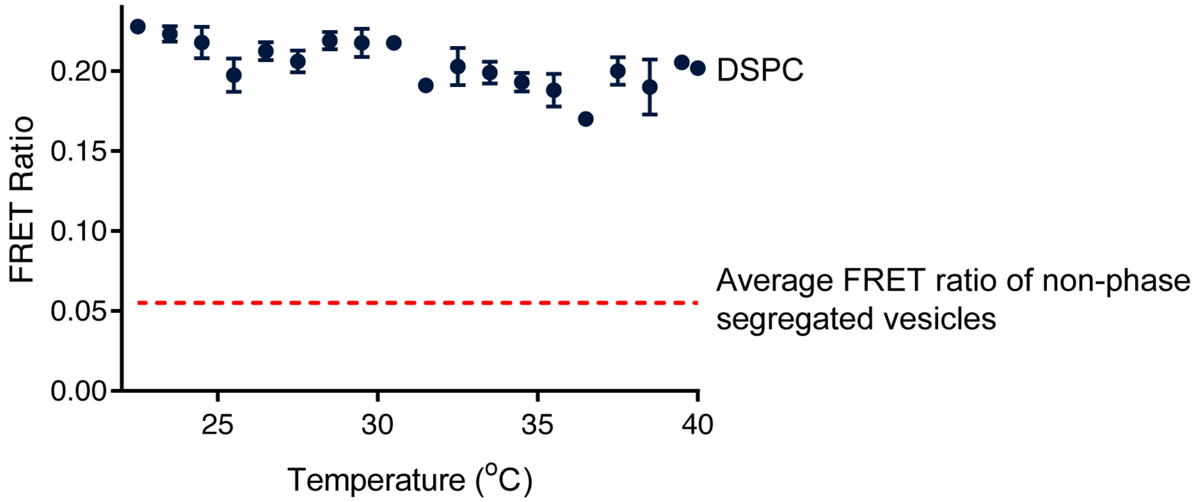


**Figure S7.** FRET ratio of 2 DSPC: 1 DOPC:1 DOPE: 1 Chol over a range of temperatures when prepared at 4 °C. The dotted red line represents an average FRET ratio for homogenous membranes composed of 3 DOPC: 1 DOPE: 1 Chol. The relatively constant FRET ratio observed over the range of temperatures tested demonstrates that phase segregation remains present at all temperatures used in this study. Error bars represent S.E.M. for a sample size of 3.

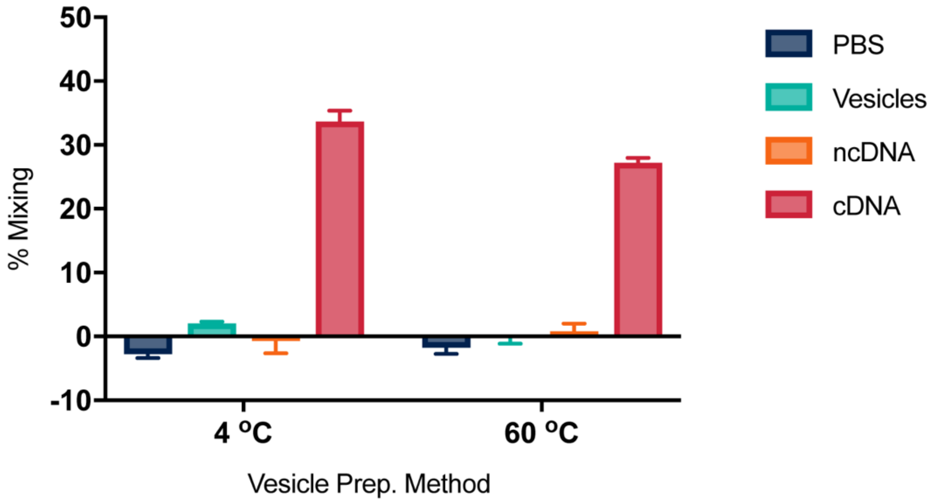


**Figure S8.** Total lipid mixing for 2 DSPC: 1 DOPC: 1 DOPE: 1 Chol vesicles prepared at 4ºC and 60ºC. This study confirms that even though the phase transition temperature for DSPC is not achieved during vesicle hydration, total lipid mixing, and thus fusion, is comparable to samples prepared at 60ºC. Error bars represent S.E.M. for a sample size of 3.

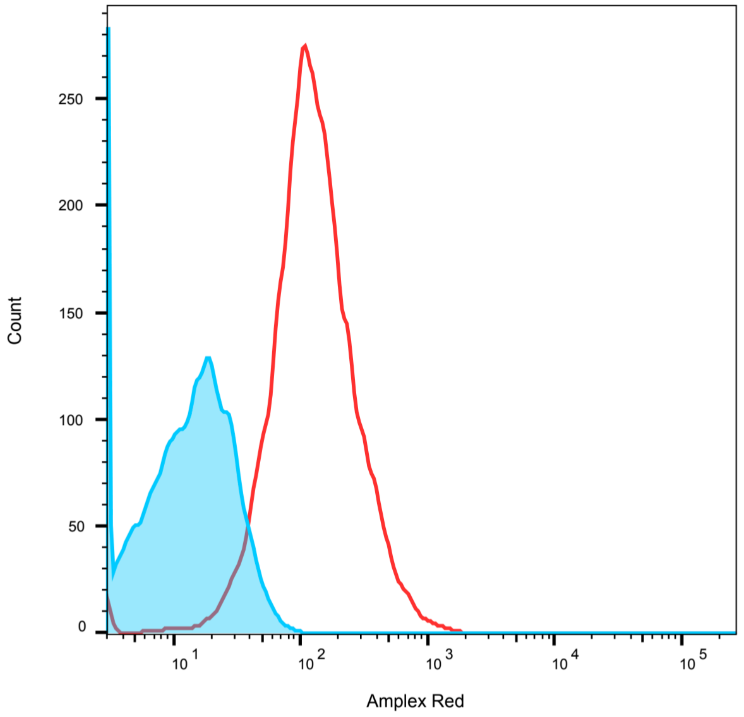


**Figure S9.** Initiation of Amplex Red reaction via DNA-mediated fusion, analyzed by flow cytometry. ncDNA vesicles fluoresce less (blue) than cDNA vesicles (red) indicating that DNA-mediated fusion allows for the progression of the Amplex Red Reaction by mixing glucose with the correct enzymes and substrates to produce resorufin. 10,000 events were recorded, p-value < 0.0001.

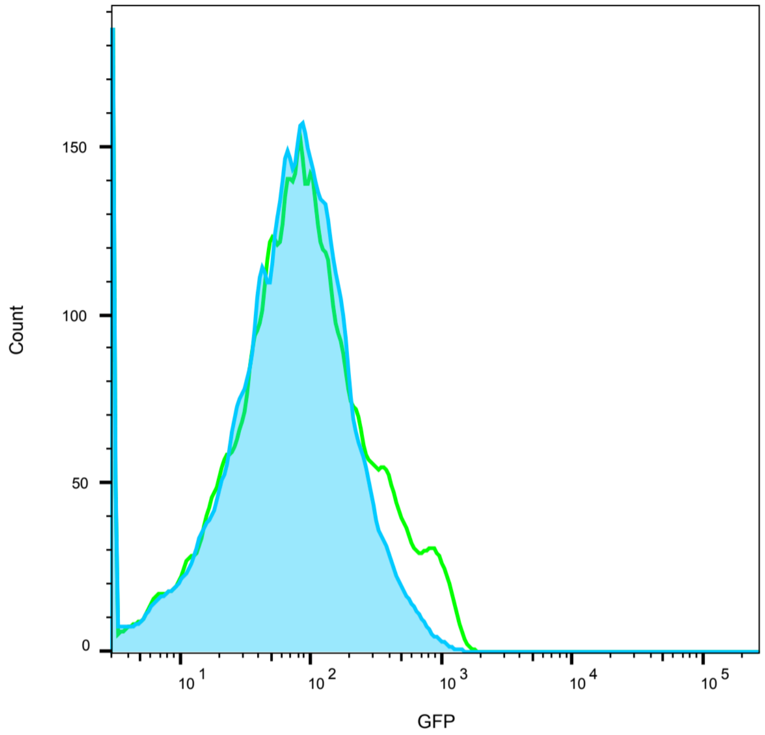


**Figure S10.** DNA-mediated fusion initiates the production of sfGFP, as analyzed via flow cytometry. ncDNA (blue) vesicles are less green shifted than cDNA vesicles. Vesicles above 1,000 nm were omitted from analysis to reduce any false GFP production as a result of autofluorescence from large vesicle aggregates. 10,000 events were recorded, p-value < 0.0001 calculated using chi-squared.
